# Reconstructing protein interactions across time using phylogeny-aware graph neural networks

**DOI:** 10.1101/2022.07.21.501014

**Authors:** David Moi, Christophe Dessimoz

## Abstract

**Motivation:** Genes which are involved in the same biological processes tend to co-evolve. Thus, metabolic pathways, protein complexes, and other kinds of protein-protein interactions can be inferred by looking for correlated patterns of gene retention and loss across the tree of life—a technique called phylogenetic profiling. Recent methodological developments on phylogenetic profiling have focused on scalability improvements to take advantage of the rapidly accumulating genomic data. However, state-of-the-art methods assume that the correlation resulting from co-evolving proteins is uniform across all species considered. This is reasonable for interactions already present at the root of the species considered, but less so for ones that emerge in more recent lineages. To address this challenge and take advantage of recent developments in deep learning methods, we introduce a phylogenetic profiling method which processes large gene co-phylogenies using neural networks.

**Results:** We show that post-processing conventional phylogenetic profiles using deep neural networks can improve predictions, but requires onerous training on specific phylogenies. Overcoming this limitation by taking the topology of the species tree as an input, Graph Neural Networks are shown to outperform all other methods when interaction detection is not centered on just one species of interest, while also predicting when interactions appeared and in which taxa they are present.

**Conclusion:** Graph Neural Networks constitute a promising new approach for phylogenetic profiling. Our work is a first foray into “dynamic phylogenetic profiling”—the reconstruction of pairwise protein interaction across time.

**Availability:** All of the code is available on the project Git at https://github.com/DessimozLab/HogProf/tree/master/pyprofiler/notebooks/Graphnet. Datasets used are hosted at http://humap2.proteincomplexes.org/download and https://string-db.org/cgi/download.

## 1 Introduction

No protein is an island. All of the complexity we can observe in biological life is due to the emergent properties of the sum of all the molecular interactions within an organism. To represent the sum total of these interactomes succinctly we can model them as protein interaction networks whose nodes represent a given protein and whose edges represent an interaction affinity. Different regions of a protein interaction network will be enriched in interactions specific to certain biological processes such as photosynthesis or signaling pathways necessary for the development of a multicellular organism. Throughout evolution, organisms speciate and adapt to new niches through incremental random changes at the gene and gene content level. Genes may be lost or become inactivated, duplicated and neofunctionalised or even gained as de-novo (Gabaldón and Koonin, 2013). These changes at the gene level, if kept, will also have an impact on the network level over time. For example, if the loss of one gene renders a particular biological process defunct, its protein interaction network neighborhood will also largely be rendered irrelevant and slowly these genes will be inactivated or lost over successive generations or repurposed for other functions in other networks.

Phylogenetic profiling methods (Pellegrini *et al*., 1999; Tabach *et al*., 2013; Franceschini *et al*., 2016; Sherill-Rofe *et al*., 2019; Moi *et al*., 2020) seek to exploit this correlated evolutionary signal between genes involved in the same biological processes in order to reconstruct the underlying pairwise interactions, interaction network neighborhoods and global interactomes.

To correlate the evolutionary histories between these gene families, a first step of reconstructing each family’s history is necessary. This can either be done by comparing the extant distribution of a homologous gene family using a simple presence/absence approach for each taxon of interest, or more complex pipelines designed to reconstruct the full history of a gene’s evolutionary trajectory from its emergence to its extant distribution. In this work, we do the latter by using as input the comprehensive gene family phylogenetic reconstructions from the well-established OMA database (Zahn-Zabal *et al*., 2020; Altenhoff *et al*., 2021).

Typically, the evolutionary information for each protein family is encoded in either a tree-like or vector object. A distance between the phylogenetic profiles representing the evolutionary signatures of protein families can be established using any number of methods appropriate for either vectors or trees. The methods that have currently been explored using vector based approaches are commonly used distances between pairs of binary or continuous vectors. Some metrics capture information on the evolutionary distance between members of a gene family in relation to the species containing the interactions that are being represented by the profiles. Using probabilistic data structures in the HOGPROF pipeline (Moi *et al*., 2020) we were able to create a fast, editable and tunable framework for profile comparison. The minhash signatures of profiles allow for the comparison and searching within sets of profiles using an approximation of the Jaccard distance (a vector based distance which had been previously established as a profile comparison metric). Other vector based distances that have been used include the Pearson correlation, empirical covariance, Euclidean distance and Hamming distance.

When using sets of closely coevolving profiles, approaches using direct coupling analysis or DCA (Morcos *et al*., 2011) may become appropriate to disentangle the conditional dependencies. Typically, in bioinformatics, these methods are used on multiple alignments (MSAs) of homologous protein sequences to find coevolving sites. These often correspond to contact points within the either tertiary or quaternary structures of the proteins which are necessary for their biological function. In a similar fashion, DCA has been applied to groups of closely coevolving profiles, either directly on the profiles themselves or on the pairs of MSAs corresponding to both families (Marmier *et al*., 2019; Fukunaga and Iwasaki, 2021). Unfortunately, however, the coevolution signal between profiles may be restricted to specific clades of species (Sherill-Rofe *et al*., 2019). Since DCA is phylogeny naive, finding the sparse inverse of the correlation matrix between profiles (within an improperly selected taxonomic subset or with a set of mismatched profiles) will result in low values with the correlation matrix. Taking its inverse will result in large eigenvalues that do not reflect the strength of interaction within a species of interest. In this work we are considering the set of all interactions and comparing metrics that can be applied to profiles which are only weakly correlated, making the application of DCA limited in its utility.

Tree-based distances are an alternative to these phylogeny naive approaches, leveraging the species tree or the phylogeny of each protein family to establish a score reflecting the coevolutionary signature of two protein families (Ta *et al*., 2011; Ruano-Rubio *et al*., 2009).

In this work, we will investigate the PPI prediction capabilities of two neural network architectures adapted to the vector and tree representation of phylogenetic profiles: deep neural network (DNN) and convolutional graph networks (CGN) respectively. To illustrate the strengths and shortcomings of each architecture we prepared two PPI datasets, one where interactions are known to be present or absent within a single species (human) and another between protein families without any restrictions on which species the interactions are found in. Designing and benchmarking methods to infer networks in the context of substantial incomplete data (Open-world assumption, e.g. Dessimoz *et al*., 2013) can be challenging, particularly when no ground truth is known that can provide true negatives for either biological functions or interactions between proteins within a subnetwork (Drew *et al*., 2021).

For the human-specific interactions we chose to use the high fidelity Hu.Map dataset (Drew *et al*., 2021). Hu.Map offers a gold standard dataset to measure our methods’ efficacy in retrieving interactions in humans, but phylogenetic profiling is also of particular interest in non-model organisms. Since ancestral interactomes diverge differentially in each clade after speciation events, interactions may be preserved in some lineages while being lost or ‘rewired’ in others (Medina *et al*., 2016). To effectively develop methods to detect these phenomena of clade specific coevolutionary signals within specific neighborhoods of the interactome, we also require a dataset spanning many eukaryotic clades. For this we chose the dataset with the broadest taxonomic range of interaction data. The STRING (Szklarczyk *et al*., 2019) database contains interaction data for protein families (co-occurring groups or COGs in their database) as well as the individual interactions contributing to the interaction score between two COGs. Using this dataset, it is possible to train classifiers to identify interactions between profiles corresponding to two COGs (in the case of the DNN) or to label a species tree to show where an interaction between two COGs may have emerged and in which extant species the interaction can be found (in the case of the CGN).

## 2 Methods

### 2.1 OMA database and HOG input data

The OMA database organizes 2500 genomes into an evolutionary data structure known as hierarchical orthologous groups—HOGs (reviewed in Zahn-Zabal *et al*., 2020). The sequence based comparison of all proteins in the OMA database is used in conjunction with the canonical NCBI taxonomic tree to derive orthology relationships for each orthogroup, the main assumption being that genes emerge at one point in the tree and are inherited by vertical descent, undergoing occasional duplications and losses.

### 2.2 Deriving evolutionary histories using pyHam

The pyHam (Train *et al*., 2018) python package allows a user to recover the evolutionary events that explain the extant distribution of the gene families contained in HOGs. Using orthology information and the NCBI taxonomic tree, pyHam outputs an annotated tree containing information on the copy number of the gene family at each taxon as well as the duplication, loss and gain events describing the gene’s evolutionary history.

### 2.3 HOGPROF-based phylogenetic profiles

The HOGPROF method efficiently identifies gene families which have similar patterns of gene gain, retention, duplication, and loss events across the tree of life (Moi *et al*., 2020). It works by encoding vectors representing the evolutionary histories extracted from HOGs by pyHam and storing these as minhash signatures (Wu *et al*., 2018). These signatures are then compiled into searchable locality sensitive hashing databases using the LSH forest algorithm (Bawa *et al*., 2005). In the context of this work, we use the explicit representation of the profile as an input for machine learning methods or vector-based distance metrics. The neural network approaches presented in this work are ideal to couple to HOGPROF as a post processing step but are too computationally intensive to exhaustively screen for interaction pairs across all HOGs.

### 2.4 Vector-based profile distance metrics

While representing phylogenetic profiles as minhash signatures with HOGPROF is useful for large-scale searching and comparison between HOGs, it may not be the best method to detect pairwise interactions or pinpoint where the interaction may be happening within the species tree. The HOGPROF method allows a user to approximate the Jaccard score between two profiles or search a profile database. The coevolutionary signal measured is an approximation of the explicitly calculated Jaccard score between two profile vectors. This signal is a good representation of the similarity between both vectors which may reflect coevolutionary signal, but direct interactions can be confounded by conditional dependencies between interaction and the score also provides no information relative to where in the profiles the coevolutionary signal is coming from. Taking into mind these limitations, using vector representations of the annotated species trees produced by pyHam we can employ strategies using the explicit vector representations of the phylogenetic profiles without calculating their minhash representation to explicitly measure profile distances or train machine learning algorithms.

Vector distance metrics that have previously been validated as relevant for describing phylogenetic profiling linkage are compared to neural network approaches in this work. Euclidean distance, Pearson correlation or Hamming distance are used alongside the explicit calculation of the Jaccard distance.

### 2.5 Deep feed forward neural network for assigning interaction probabilities

Establishing a function to classify objects—in this case pairs of profiles—into categories of interacting vs. non-interacting pairs is a classic machine learning problem. Our input data can either take the form of trees or vectors as discussed previously. In this work we explore neural network architectures that are able to solve this classification problem in order to compare them to classical distance metric based approaches. Each representation of the data has its own associated architecture which is most appropriate to take on a classification task. In the case of vector representation, it is fairly straightforward to see that a deep feed forward neural network is an appropriate architecture. This net can be trained to output a continuous value representing the probability of an interaction given the data. By training this type of architecture, we fix the input size of the net at its creation.

By training a DNN to output a prediction score between 0 and 1 on a set of positive and negative pairwise interactions, we can use it to predict the probability of interaction based on any two profiles for proteins that could be participating in the set of interactomes used to train the network.

The network will learn to output predictions on the probability of a given PPI based on pairs of evolutionary histories for one particular clade or organism of interest. Deciding the taxonomic composition of the PPI pairs included in the training set thus presents a tradeoff between specificity and versatility. Also, as new genomes are sequenced and their orthology relationships are derived, the input set of species (and taxa in the species tree) will change. This change in the input requires the network to be retrained to use this approach to predict PPIs.

In this work, we implement a fully connected feed-forward network architecture with 21048 input units corresponding to the number of possible loss and duplication events or presence for two labeled taxonomies corresponding to two profiles), 200 hidden units in the first layer and 100 hidden units in the second layer with sigmoid activation functions feeding into a final layer with one unit. This network is implemented using Tensorflow (Abadi *et al*., 2016) and Keras (Chollet, 2015). The network was trained on the binary vector representation of the phylogenetic profiles constructed with HOGPROF as detailed in Sect. 2.3 to predict a binary output of interaction presence or absence between two HOGs. Preparation of Training and testing sets with STRING and Hu.Map is explained in Sect. 2.8. Full parameters for both trained models are available in the git repository associated with this manuscript.

### 2.6 Reconstructing the evolutionary history of an interaction pair using the Fitch parsimony algorithm

The physical interaction between two protein sequences can be thought of as a phylogenetically inherited trait. Across extant species we can perform experiments to derive which sequences are in interaction. By combining this interaction data with orthology information, linking each of these interactors to a protein family, we can derive in which species two families have experimental evidence of interaction. By assuming vertical inheritance of this trait along the species tree, we can reconstruct the ancestral binary states representing the presence or absence of this interaction using the Fitch algorithm (Fitch, 1971). This approach presupposes that the most parsimonious explanation for the extant distribution of interactions is the correct one. For the purposes of training graph neural networks to label species trees with binary interaction states, we will assume that our Fitch reconstruction of interaction state is the ground truth.

### 2.7 Convolutional graph neural networks and reformulation of PPI detection as a graph labeling problem

Phylogenetic trees can be represented as graphs and the problem of classifying pairs of profiles into interacting or non-interacting categories can be turned into a graph labeling task. Fortunately, neural network architectures have been designed for precisely this type of task. One such architecture is a CGN (Li *et al*., 2015), which can be used to either classify entire graphs or label nodes within a graph. As orthology datasets expand and more species are included in phylogenetic profiles, it is indeed useful to have a network architecture that can adapt to new tree topologies without needing to be retrained for a new input set. Also, training a DNN to recognize interacting profile pairs which correspond to interactions within one species is a fairly straightforward task, with evolutionary events present indicating this interaction confined to the same branches of the tree of life. These should appear across many samples of the dataset and provide a reliable signal for the DNN to classify profile pairs. However, protein families may only be interacting within a subset of species of the species tree. A DNN architecture will generalize poorly to this type of problem since features which may be informative for finding interaction in one species will be totally irrelevant for another. In contrast, a CGN will be able to generalize to label all species using information about the structure of the species tree graph and the events that happened leading to each node.

While explaining the principles behind graph neural networks is beyond the scope of this work, it can be useful to think of them in similar terms as recurrent neural networks (which could be thought of as a subset of CGNs designed to work with a graph where nodes are only connected to the nodes representing the previous and next points in a given sequence). Each node receives a set of messages from its neighbors and performs a transformation on a permutation invariant aggregation of the messages. This confers a learned context dependent embedding to each node which can be used by downstream functions to infer outputs such as node or graph labels.

In our use case, by representing the species tree as a directed graph, profiles can be reformulated as annotated graph objects by storing the evolutionary information derived with pyham in each node. Additional information related to the topology of the species tree relative to each specific node (e.g. the number of child nodes or sister nodes) can be added to the node annotation as well. Now, instead of comparing two trees with a vector or tree based metric we define the problem as a graph labeling exercise, either labeling the entire aggregated graph as indicative of global coevolution between two families, or locally at each node, indicating regions which show signs of coevolutionary signal. Since the species tree is a directed acyclic graph, we can exploit this property of directionality in a directed graph, creating two layers of taxonomic nodes, one for passing information up the graph and another for passing it downward.

The CGN network presented in this work is implemented in the pytorch geometric package (Fey and Lenssen, 2019). A convolutional graph network architecture was used with 3 layers comprising 50 hidden units each connected to an output layer using a *tanh* activation and a bias term. The convolution layers first used in (Duvenaud *et al*., 2015) were used for the convolution layers for the phylogenetic graph to allow for the adaptive weighting of evolutionary events relative to their distance to a target node. A transformer convolution layer (Shi *et al*., 2020) was used to input all of the HOG nodes to a single node representing the overall state of the graph (global presence or absence of interaction) to make use of an attention mechanism. The choice of using an attention mechanism was made to selectively weigh informative HOG nodes as input for this prediction.

This net was trained using STRING and Hu.Map data separately. An early stopping point was manually selected once the net stopped showing improvement over 20 epochs. Full parameters for the construction and training of the net are available in the git repository associated with this manuscript.

### 2.8 PPI data preparation

The Hu.Map dataset contains a total of 17,526,311 PPIs with an associated interaction score based on different types of evidence. The PPIs were filtered to those with a score above 0.75 probability resulting in 8360 interaction pairs. These were considered True positives. Then, by shuffling the interactors of these pairs randomly, a set of True Negatives was generated. The data was split between training and testing sets with a .7 and .3 split with equal numbers of positive and negative samples.

All of the proteins in the dataset were mapped to OMA HOGs (April 2021 version) and tree and vector profiles were generated with HOGPROF to be used with the profiling strategies presented in this work.

The STRING database has its own implementation of orthologous groups called Clusters of Orthologous Groups (COGs) (Szklarczyk *et al*., 2019). An interaction score between COGs comprising the sum of coexpression, experimental, database and text mining evidence channel scores was calculated for all COG interaction pairs. COG pairs with an interaction score above 1000 were selected as a first true positive dataset. A first set of Negatives for hyperparameter refinement were generated by shuffling the initial set of True positives as with the Hu.Map dataset. An optimal filtering of the dataset was found by observing the effect of additional cutoffs for each evidence channel on the dataset size and by using jaccard distance to derive an AUC for detecting interaction. An additional threshold of a score of 500 for the text mining evidence channel was selected due to its beneficial effect on AUC while maintaining a sufficiently large dataset for training. These pairs were considered the final set of True positives for the STRING dataset. Then, by shuffling this final set of True positive COGs interactions randomly, a final set of True Negatives was generated. Each COG was then mapped to its individual protein identifiers within STRING. The STRING dataset contains a total of 20 billion PPIs between proteins annotated with multiple evidence channels. To determine in which species there was evidence for each interacting COG pair, we checked for protein pairs within the same species from both COGs for all COG pairs within our filtered dataset. Since looking up each pair would have been prohibitively expensive computationally, a set of Bloom filters (with a false positive rate of 0.01 for 10^9^ elements) was established containing the signatures of all protein pairs within our COGs of interest in STRING. After checking for the presence of interaction pairs in the intersection of the set of species found in both COGs using the Bloom filters, a parsimony-based reconstruction of interaction along the taxonomy was calculated for each COG pair as described in methods 2.6. In figure 2 below, a single sample of two COGs that have interaction data in some species in STRING is shown.

**Fig. 1.**
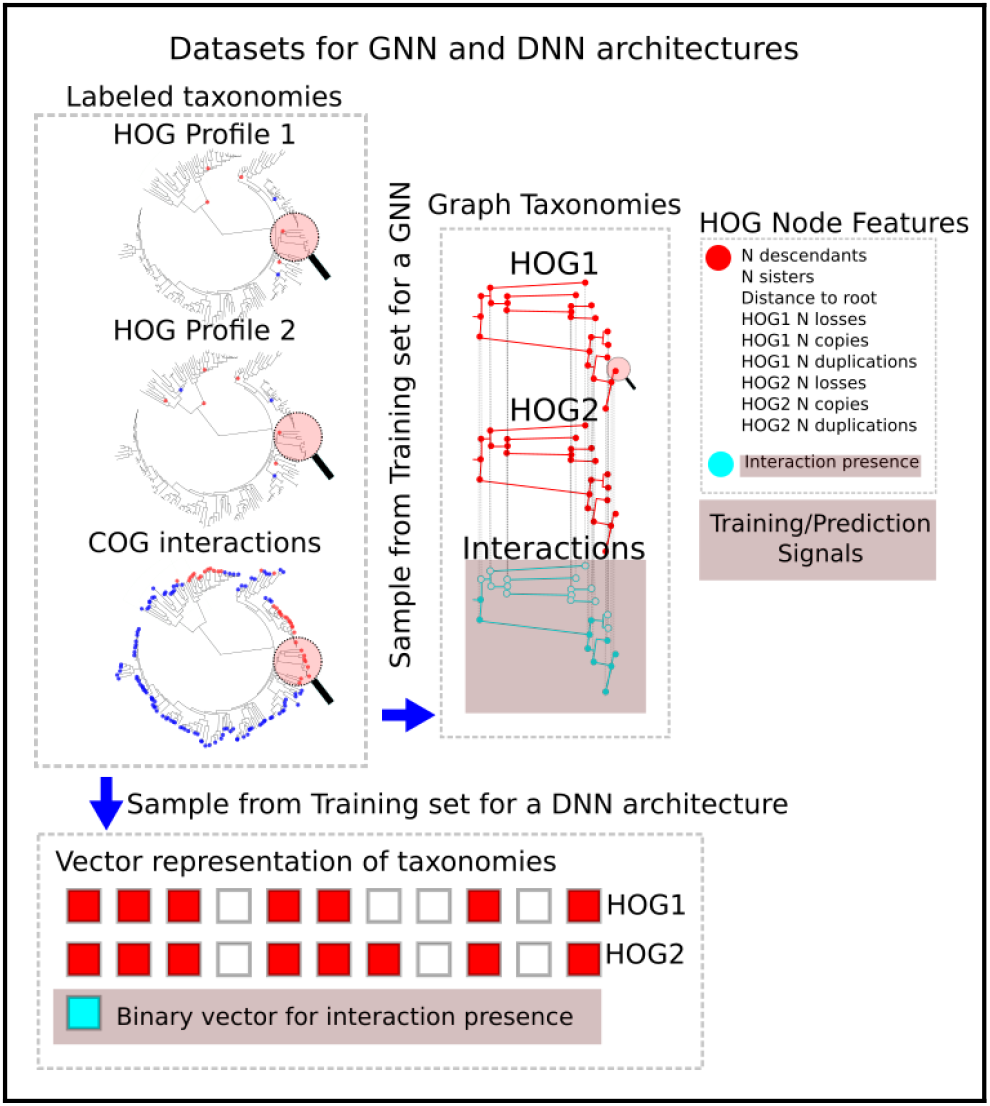
Pipeline for labeling pairs of tree profiles with graph neural networks. a) Species trees are annotated using STRING PPI data showing evidence of an interaction between two HOGs in a species. The interactions are then propagated up the tree to reconstruct the ancestral binary state of the interactions using parsimony. Here the leaves are shown with blue and red dots to depict a hypothetical case where interactions are either present or absent in the STRING database between two HOGs in each species of the taxonomy. The HOGs are used with pyHam to create annotated species trees showing the evolutionary events in the family’s history. b) Taxonomies are transformed to graphs. The convolutional graph neural network is trained to annotate a species tree graph. The pyham labeled taxonomies provide information for each node of the and the reconstructed interaction states are the node labels to be inferred. c) Each node has the pyHam information on nodes of the species trees representing the evolution of pairs of HOGs. Pairs that have no evidence of interaction have empty species trees with no positive labels for interaction whereas trees with evidence have a parsimony based reconstruction of interaction states. d) The DNNs are trained on a binary vector representing presence, loss and duplication on each branch of the taxonomy. The net is trained to perform a binary classification task to decide whether or not the pair of profiles are interactors.

**Fig. 2.**
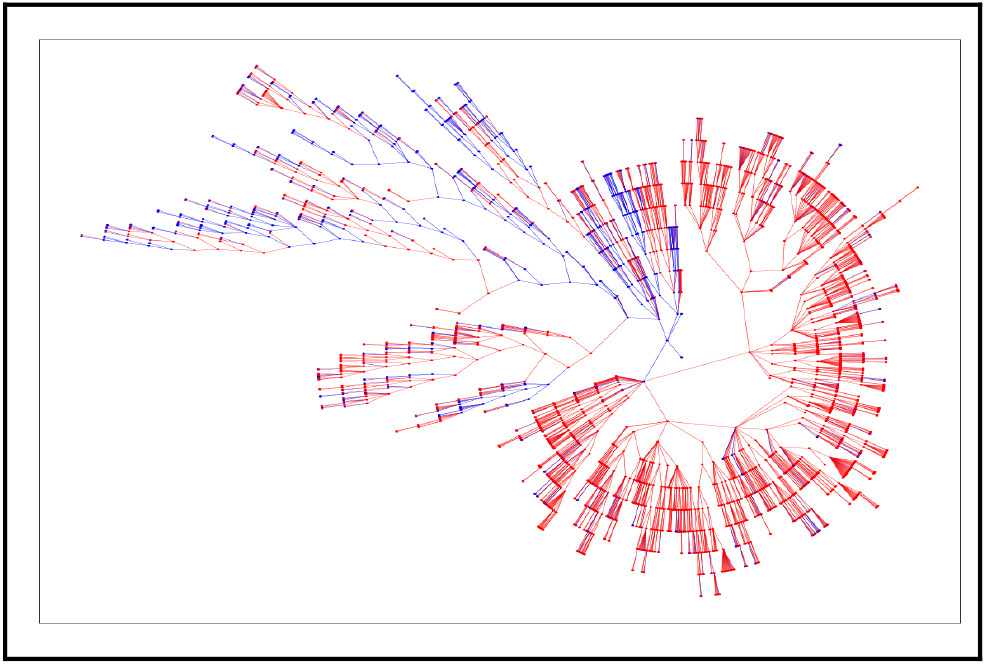
An example of a positive phylogeny graph sample from the STRING dataset. After checking for interaction presence between COG0088 and COG1756 in all species present in STRING within the bloom filters, the leaves present in the OMA taxonomy are labeled with either 1 or 0 (interaction presence or absence respectively). In this graph blue denotes the presence of an interaction and red denotes the absence. The Fitch algorithm is then used to generate a reconstruction of the ancestral interaction states. In this graph visualization of the taxonomy, the node distance from the root of the tree is represented as the radius. Each branch length is set to 1 since the NCBI taxonomy does not contain branch length information.

For samples of non-interacting pairs, the ground truth taxonomy was labeled as non-interacting in all nodes in the intersection of the set of species where both COGs are found. The data was split between training and testing set with a 0.7 and 0.3 split with equal numbers of positive and negative samples. All of the COGs in the dataset were mapped to OMA HOGs (April 2021 version) and profiles were generated with HOGPROF (methods 2.2).

While the STRING database incorporates a wide breadth of species and is ideal for this use case requiring interaction data across all of the leaves of the NCBI taxonomy, it is important to keep in mind its limitations and biases. Data for 14,094 organisms are found in the dataset, but the bulk of interaction data corresponds to heavily studied eukaryotic (and to a lesser degree prokaryotic) model species. This degree to which certain clades of the tree of life are heavily studied or ignored in PPI studies will be reflected in the completeness of the data and lead to false negatives in clades where interaction data is sparse. This will in turn impact our parsimony based reconstruction of the ancestral states. Additionally, evidence channels in the description of the evidence for each interaction may regroup several methods (e.g. experimental) which may not be comparable or equally reliable in different clades of the tree of life.

## 3 Results

### 3.1 Interaction prediction using Hu.Map data

Both CGN and DNN network architectures were trained on STRING and Hu.Map Training datasets as detailed in *Methods*. As interaction data was not available in all species in Hu.Map only a pairwise interaction probability between two profiles was inferred using this dataset. The two net architectures were used to infer an interaction score metric between profiles corresponding to two STRING COGs or two Hu.map entries alongside vector based distance metrics. In the use case of learning to predict interaction in a single species, the DNN solidly outperforms phylogeny explicit distance metrics in the Hu.Map dataset. This is shown below in figure 3.

**Fig. 3.**
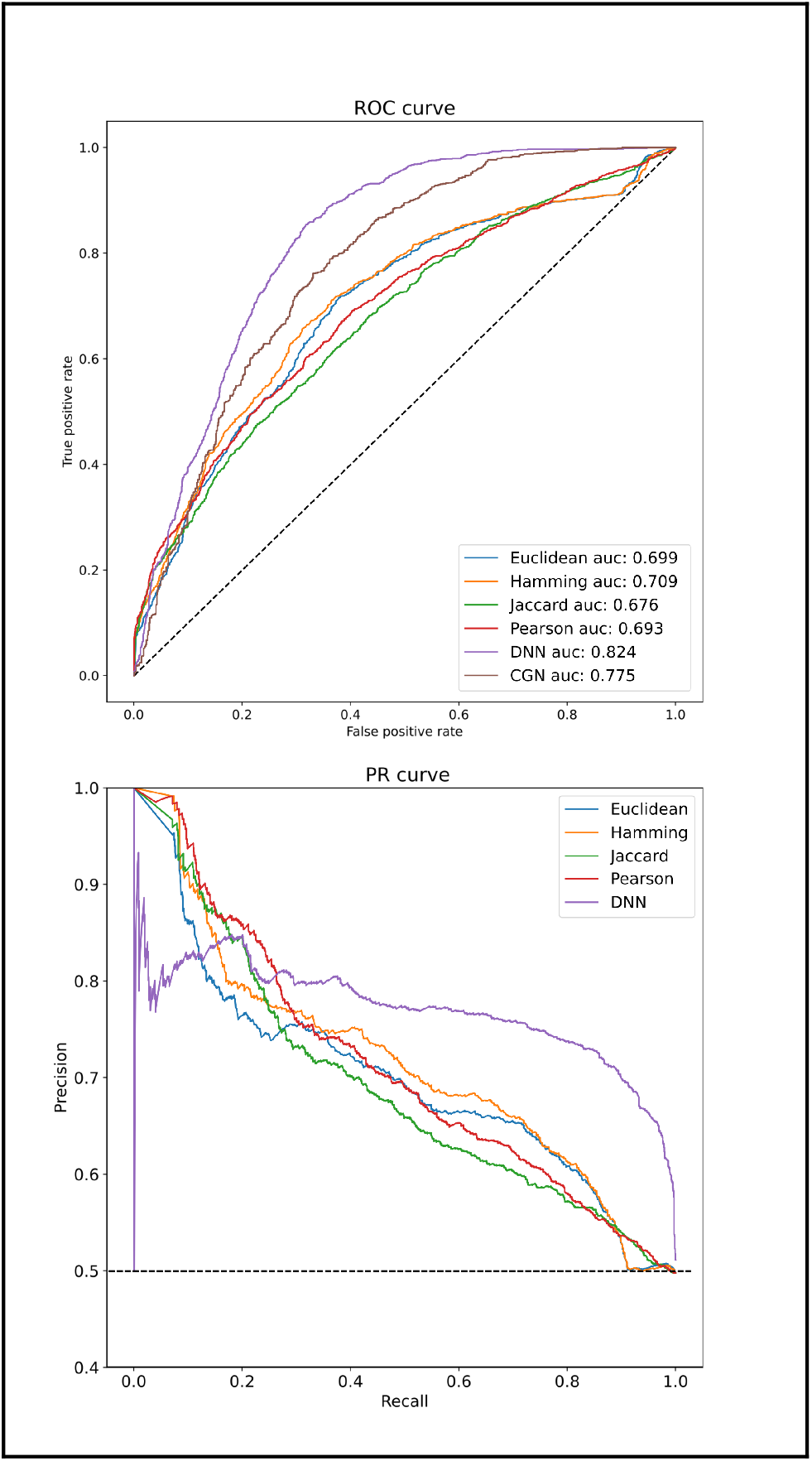
Graph level prediction using Hu.Map data. The ROC curve for the DNN based classifier shows the best performance in terms of AUC characteristics when compared to simple distance metrics when trained. Interestingly, if high precision is required, explicit distance metrics appear to perform better than neural network approaches. Considering the ease with which these can be calculated, they could potentially be included as inputs to a neural network approach in tandem with the profile features, potentially remedying this issue as the network learns to integrate this signal.

### 3.2 Interaction prediction using STRING data

The results using the CGN on this dataset show that, despite lacking awareness of taxonomic nodes’ identities and only having information on graph topology and evolutionary events, it is able to detect coevolutionary signal between profiles at a higher AUC than vector distance metrics. While all approaches can provide a score for the global probability of interaction, the real interest of the CGN lies in its graph labeling capacities. It is also able to identify clades in which the interactions are likely to be happening. In figure 5, these predictions are shown in the plot under the title ‘taxon level prediction’.

While it may be appropriate for a use case where the interaction detection is centered in a species or clade of interest, the DNN’s prediction ability suffers when interactions are not always inferred in the same clades. However, the permutation invariant character of CGNs is ideal for dealing with this challenge. This difference in prediction quality is reflected in the ROC curves for STRING DNN and CGN architectures shown below in figure 4. The DNN underperforms even explicit distance based metrics due to overfitting the training set (despite using dropout (Srivastava *et al*., 2014) during training) whereas the CGN provides a reliable signal of global interaction as seen below in figure 4.

**Fig. 4.**
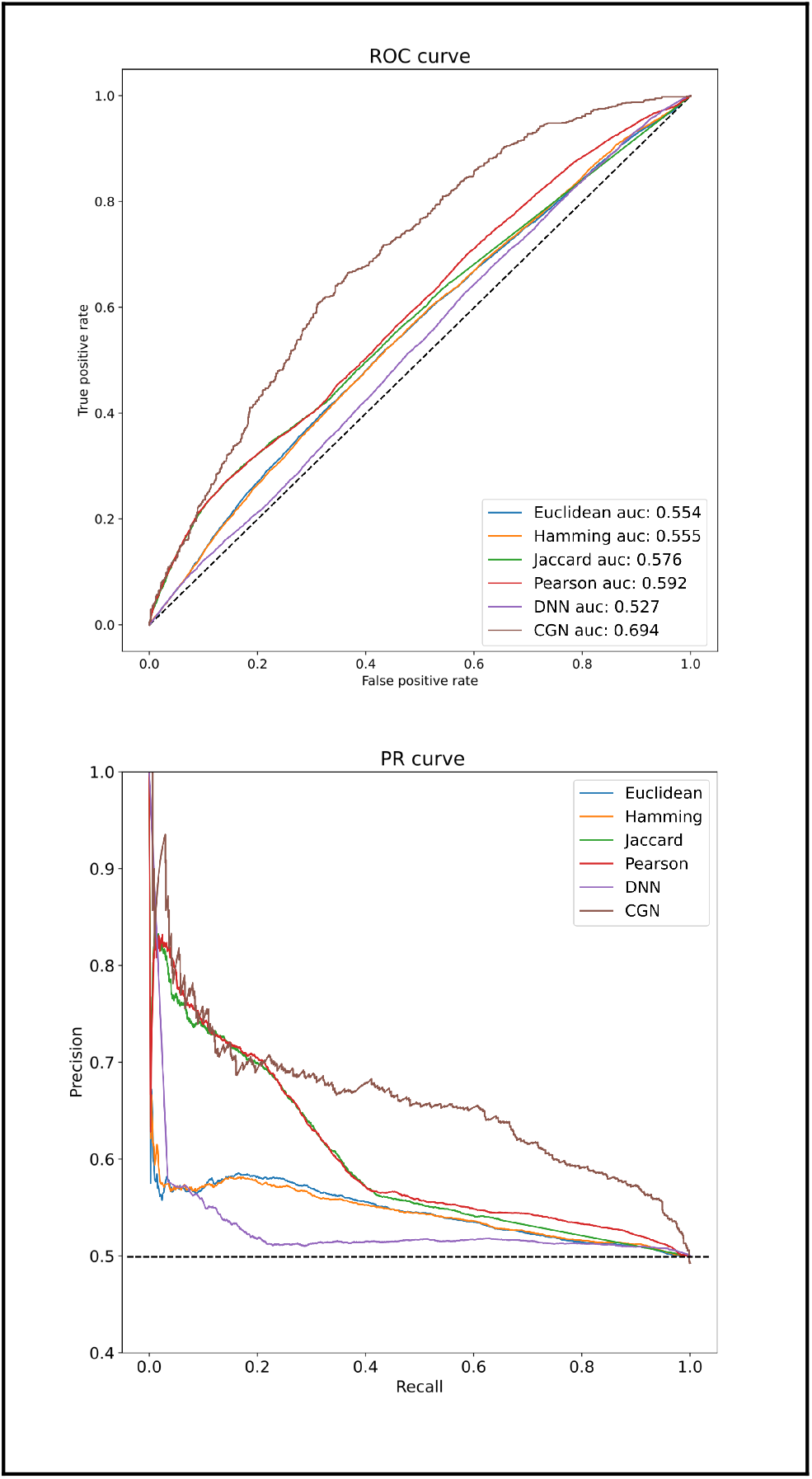
Graph level prediction using STRING data. The DNN is unable to converge to a consistent pattern due to the shifting point of reference of the species of interest in which interactions are found. The explicit distance metrics perform worse than in the case with Hu.Map data pointing to a change in the separability of the positive and negative samples in this dataset relative to the high fidelity human interaction set. The CGN architecture outperforms all approaches in this use case and is able to extract informative signals despite the noisy nature of the dataset.

In addition to providing a way to circumvent the permutation of the point of reference for interactions within the graph, the CGN architecture also allows for the user to predict interaction on individual graph nodes. As far as we are aware, this is the first effort that has been made using profiling data to reconstruct interaction history over the taxonomy using phylogenetic profiles coupled to an artificial intelligence approach, although some other approaches to reconstruction ancestral interaction states have been proposed using PPI data as input such as (Rajan *et al*., 2021), for example. Below in figure 5, the performance of the net’s predictions compared to the ancestral reconstructions detailed in methods 2.6 is shown. While the quality and completeness of the data present in STRING is highly correlated to the clade containing the interaction data, we nevertheless chose this database for its breadth in order to ensure a maximum coverage of all of the species present in OMA based phylogenetic profiles. In addition, OMA profiles are also based on the NCBI taxonomy, which is limited in its topological information due to polytomies and lack of branch lengths. With the limitations of the graph dataset we constructed in mind, it is still exciting to note that the net is able to provide predictions for each node solely from the signal present in the profiles of pairs of HOGs.

**Fig. 5.**
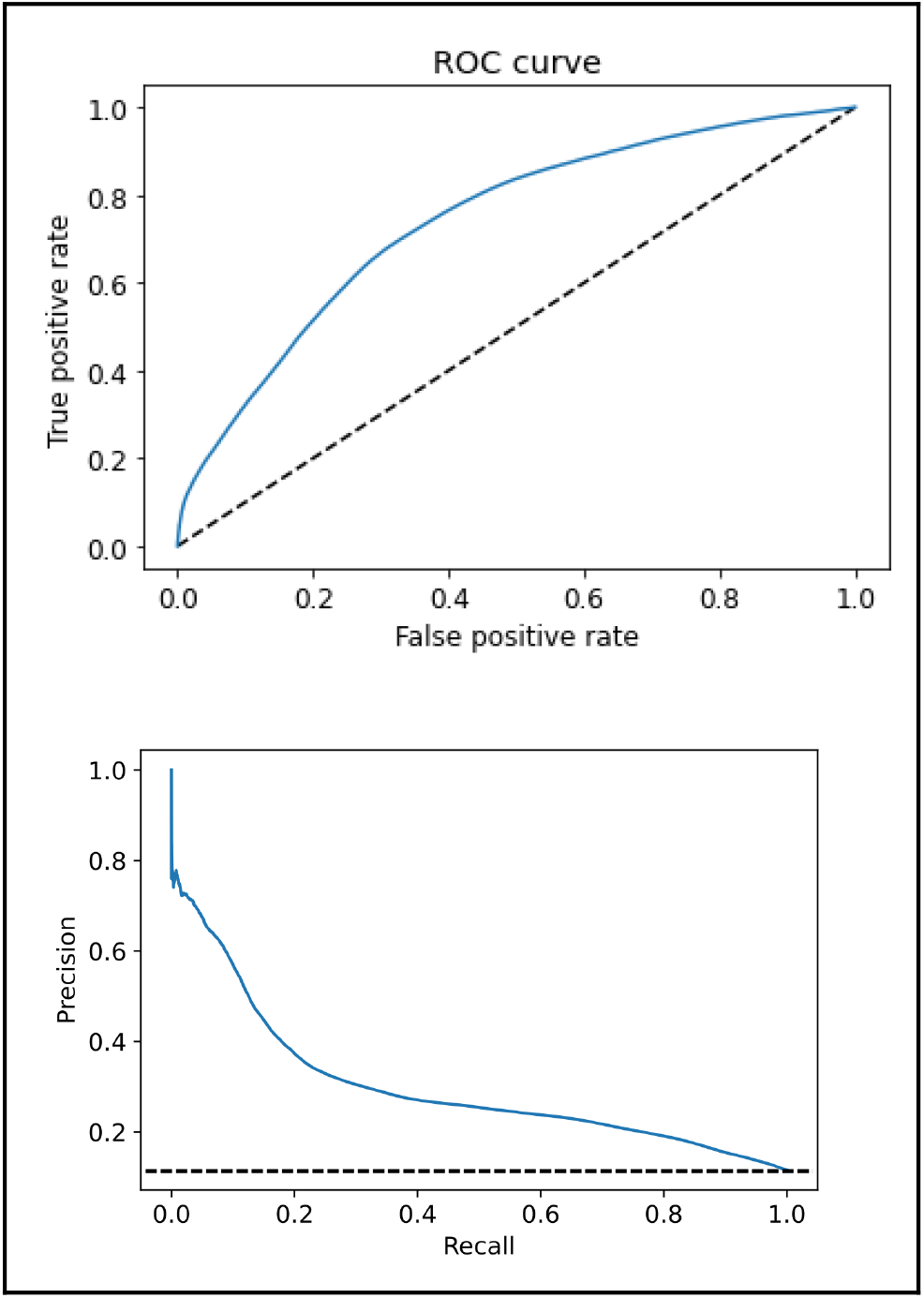
Taxon level interaction prediction for STRING dataset. The CGN is the only method capable of assigning an interaction probability to each node of the taxonomic tree. In this ROC curve we see the CGN’s ability to reconstruct interaction histories in both the positive and negative samples (where the states are all set to 0). However, the data the net is trained on is a parsimony based reconstruction of incomplete and noisy data with a compositional bias (as mentioned in methods 2.8). Despite these limitations, the net does find signal and is able to assign clade specific interaction probabilities between HOGs and obtains an AUC of 0.74 against the test set.

## 4 Discussion

In this work we explored the possibility of using phylogenetic profiles as input to neural networks with CGN and DNN architectures. Using only a toy model with a DNN approach, it is possible to extract phylogenetic features from the gain, loss, duplication and retention patterns that are relevant to reliably predict protein-protein interactions within one species. These models are lightweight to train and deploy and could be imagined as a potential filter in online profiling approaches such as the one currently implemented in the OMA browser. However, these features are not permutation invariant with regards to the graph structure of the phylogeny and can only provide predictions for one species or a small clade reliably.

As seen with our example using interactions between string COGs, by changing the species or set of species in which our interactions may be taking place, we cannot learn features that are consistent across the dataset to allow us to predict interactions in arbitrary clades with DNNs. By using a CGN architecture we circumvented this problem, creating embeddings of each node within our graph object representing a pair of annotated taxonomies. Each of these nodes can be labeled using its graph neighborhood rather than a fixed set of features, allowing for the prediction of interaction across all taxa in the tree of life with a single model. The speed of prediction is also comparable to a DNN approach and can also be served online in a similar context, allowing a user to screen for potential interactions within a clade or species of interest based on orthology data alone. This approach provides a guide to experimentalists looking for a starting point in describing the evolution of an interaction network in a species of interest or across the entire tree of life and can complement other sources of interaction data. In future work we plan on augmenting this approach using higher quality trees including branch length information and incorporating multiple sources of interaction data.

In addition, representing phylogenies as graphs may aid in the detection of residue level coevolution such as detecting contacts in protein structures from a sequence of residue level transitions in two columns of an alignment associated to a phylogenetic tree. This could serve to complement approaches such as DCA or serve as input data to other structure prediction methods. Aside from phylogenies, many other biological datasets or objects are easily transformable to graph representations and could benefit from the node or graph classification as well as the link prediction task CGN architectures lend themselves well to. Efforts to apply them have already seen success using protein structures (Gligorijević *et al*., 2021), PPI and coexpression networks (Xiao and Deng, 2020), metabolic networks (Harada *et al*., 2020) and they may even be used at broader scales in future projects with ecological networks (Guo *et al*., 2020). It is also tempting to imagine multigraphs incorporating a combination of objects and links mixing phylogenetic, interaction and structural graphs to develop more holistic embeddings benefiting from orthogonal sources of information.

We will continue experimenting with novel graph and network architectures and hope that this first exploration will lead to further work using phylogenetic graphs with AI.

## Acknowledgements

Many thanks to my amazing labmates, Pablo Aguilar for his help in revising this manuscript and Rocio Ballesteros for our interesting discussions on machine learning methods related to the topic of coevolution.

